# Expanding the Galaxy’s reference data

**DOI:** 10.1101/2020.10.09.327114

**Authors:** Nagampalli VijayKrishna, Jayadev Joshi, Nate Coraor, Jennifer Hillman-Jackson, Dave Bouvier, Marius van den Beek, Ignacio Eguinoa, Frederik Coppens, Sergey Golitsynskiy, Michał Stolarczyk, Nathan C. Sheffield, Simon Gladman, Gianmauro Cuccuru, Björn Grüning, Nicola Soranzo, Helena Rasche, Bradley W. Langhorst, Matthias Bernt, Dan Fornika, David Anderson de Lima Morais, Michel Barrette, Peter van Heusden, Mauro Petrillo, Antonio Puertas-Gallardo, Alex Patak, Hans-Rudolf Hotz, Daniel Blankenberg

## Abstract

**Summary:** Properly and effectively managing reference datasets is an important task for many bioinformatics analyses. Refgenie is a reference asset management system that allows to easily organize, retrieve, and share such datasets. Here, we describe the integration of refgenie into the Galaxy platform. Server administrators are able to configure Galaxy to make use of reference datasets made available on a refgenie instance. Additionally, a Galaxy Data Manager tool has been developed to provide a graphical interface to refgenie’s remote reference retrieval functionality. A large collection of reference datasets has also been made available using the CVMFS repository from GalaxyProject.org, with mirrors across the United States, Canada, Europe, and Australia, enabling easy use outside of Galaxy.

**Availability and implementation:** The ability of Galaxy to use refgenie assets was added to the core Galaxy framework in version 20.05, which is available from https://github.com/galaxyproject/galaxy under the Academic Free License version 3.0. The refgenie Data Manager tool can be installed via the Galaxy ToolShed, with source code managed at https://github.com/BlankenbergLab/galaxy-tools-blankenberg/tree/main/data_managers/data_manager_refgenie_pull and released using an MIT license.

## Introduction

Among the primary resources required when performing biomedical genomic analyses are the collection of reference datasets and annotations, including sequences and features (such as genes, locations of epigenetic modifications, regulatory regions, etc.) as well as derived datasets such as index files for sequencing-read mapping tools e.g. Bowtie 2^1^ or BWA^2^, FASTA sequence indexes with SAMtools^3^, etc. While generating the derived data is relatively straightforward, such as running “bwa index -a bwtsw reference.fa” or “samtools faidx reference.fa”, these important reference data have to be managed efficiently. Moreover, in order to allow for reproducible analyses, not only do genome build versions (e.g. hg19 vs. hg38) tool versions (e.g. indexed with BWA version 0.7.17-r1188) need to be recorded, also algorithmic choices (e.g. algorithm for constructing BWT index: “bwtsw” or “is”) are essential. This has led to the development of several software solutions for reference genome resource management.

We have previously developed the Galaxy Data Manager framework^4^ which enables provenance-backed and reproducible reference datasets inside of Galaxy^5,6^. The Data Manager framework leverages the Galaxy tool system to enable community-driven creation and dissemination of reference dataset build recipes as Galaxy tools. These Data Manager tools can be installed on-demand from the ToolShed^7^ to allow Galaxy administrators to fetch, build, and install new reference data. All underlying tool dependencies (SAMtools, BWA, STAR^8^, Kraken^9^, etc.) are well defined with versions and can be automatically and reproducibly resolved using e.g. Bioconda^10^ and Docker. As with any Galaxy tool, multiple versions of a Data Manager can be installed and executed, with provenance and reproducibility maintained. Data Manager tools can be accessed using the Galaxy GUI, a RESTful API, or by using command-line scripts. Currently, over 70 Data Manager tools have been created and shared on the ToolShed by the Galaxy community.

Refgenie^11^ has been recently published as a reference genome resource manager. Refgenie provides a command-line interface to download, build, and access reference genome assembly assets (i.e. reference sequences, mapping indexes, etc.). It can also be controlled via an API from other software. There are currently 24 recipes available for building assets locally. Refgenie can also provide a lightweight web server enabling sharing of local resources.

In light of its potential to become a widely accepted standard for reference data management, we have enabled the use of refgenie-based assets inside of Galaxy. Users are able to seamlessly access refgenie-defined references within Galaxy tools, and administrators are able to access refgenie’s ability to download reference datasets from remote resources. This approach enables the simultaneous creation and use of reference files by both Data Managers and refgenie within Galaxy.

Moreover, the adoption of refgenie by the Galaxy community will make high quality reference data sets and derived assets available for the broad scientific community.

## Methods

We have extended the Galaxy Data Manager framework to interoperate with refgenie, a standalone reference genome manager.

### Using refgenie assets in Galaxy

Integration of refgenie with Galaxy is available as of Galaxy version 20.05. Enabling refgenie usage within Galaxy is accomplished by following an optional two-step process: installing refgenie and configuring Galaxy.

A server administrator must first install refgenie using the standard prescribed instructions and take note of the path selected for its genome configuration file (see also https://galaxyproject.org/admin/refgenie/ and supplementary materials).

Second, refgenie is enabled in Galaxy by setting the ‘refgenie_config_file’ value to the previously chosen genome configuration file path within the primary Galaxy configuration file (e.g. ‘galaxy.yml’; figure 1A). This setting enables the default mappings between refgenie asset names and values into Galaxy data table entries. Currently, default mappings have been defined for genome build identifiers, chromosome sizes and sequences, SAMtools FASTA indexes, Bowtie 2 indexes, BWA indexes, and HISAT2 indexes. Additional mappings can be defined as shown in figure 1B. The assets loaded from refgenie, are merged with those from Galaxy’s native Data Managers. This approach provides maximum flexibility for the administrator while creating a single unified access point from the Galaxy user’s perspective. In fact, it is possible for an administrator to define multiple different refgenie configurations within a single Galaxy instance, making the assets defined in each resource available as a combined reference collection in Galaxy. This can be helpful, for example, if the administrator wants to provide access to the prebuilt collection of datasets in CVMFS (see below) along with additional locally maintained refgenie datasets.

**Figure 1.**
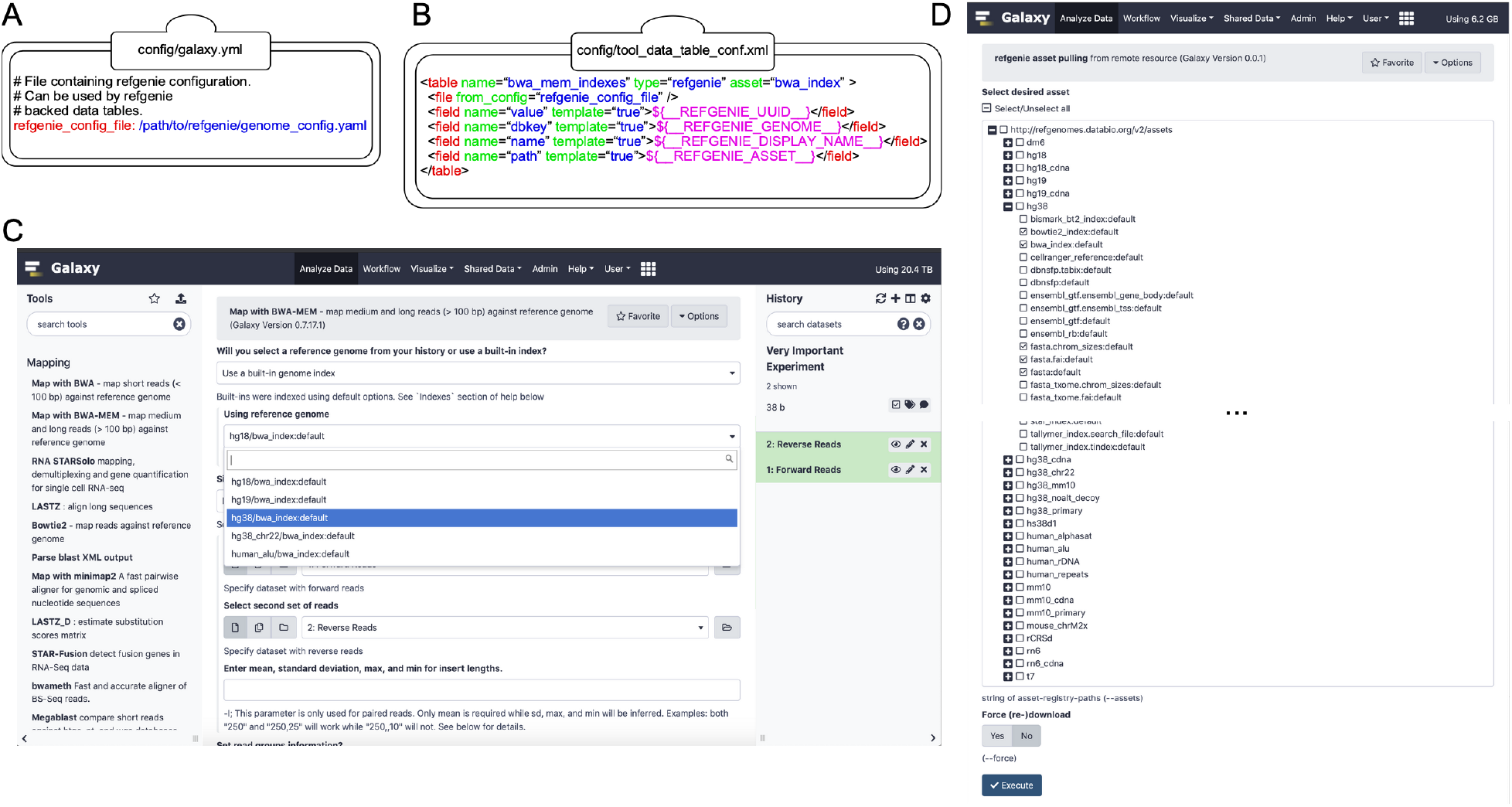
Extending Galaxy’s reference data with refgenie. (A) Setting the value of ‘refgenie_config_file’ to the previously chosen genome configuration file path within the primary Galaxy configuration file (e.g. ‘galaxy.yml’). (B) Example data table mapping between refgenie assets and Galaxy data tables for the BWA tool. Cheetah templating language is used to specify mappings between values, with several pre-populated refgenie variables available as shown. (C) refgenie assets are available for users to select and use in the Galaxy BWA tool. In this example, the user is mapping a set of paired-end sequencing reads against the hg38 genome. (D) A dynamically generated list of available remote refgenie assets are listed for an administrator to select in the ‘refgenie pull’ Galaxy Data Manager tool.

Once the refgenie integration has been configured, the users of the Galaxy server will be able to select the refgenie-provided datasets in Galaxy tools that make use of the data tables for which a mapping is defined (figure 1C).

### Retrieving additional refgenie assets in Galaxy

An initial refgenie installation contains no data. However, the administrator is able to search, download, and install assets into refgenie from public remote servers. Any asset loaded into refgenie using the standard approaches will be made available within Galaxy. To facilitate management of the assets, a Galaxy Data Manager tool has been created to provide a graphical interface to refgenie, which is now available in the Galaxy ToolShed^7^. When a Galaxy administrator accesses this Data Manager, a list of available remote assets is dynamically populated and displayed (figure 1D). These available references are organized by remote resource, genome build, asset type, and tag. The administrator selects one or more assets to retrieve, and then clicks “Execute”. An external process is queued, using Galaxy’s job management system, which calls refgenie’s standard command-line tools to perform the requested actions. Once refgenie has completed, Galaxy loads the new values and makes them available to users. This Data Manager can also be controlled using the standard Galaxy tool API.

### A CVMFS mirror

The refgenie software provides reference asset downloads from http://refgenomes.databio.org/. To increase reliability and availability, we have created a mirror of all of Galaxy and refgenie reference data available in a CernVM-FS^12^ (CVMFS) repository. CVMFS is a caching, HTTP-based filesystem with a Filesystem in Userspace (FUSE) (mount) client. The Galaxy CVMFS repository distributes the data in multiple replicas across the United States, Canada, Europe, and Australia. Moreover, it is possible to make the mirrored data directly available on any other server (see https://galaxyproject.org/admin/reference-data-repo/). When initially mounted, CVMFS does not consume any additional local disk space. Instead, as files are accessed, they are pulled from one of the replica (Stratum 1) servers to a local disk-based cache of a configurable size. This allows the administrator to directly load the refgenie configuration from e.g. /cvmfs/refgenomes-databio.galaxyproject.org/genomes_config.yaml, without needing to predetermine which assets the users may desire.

The presented approach of data distribution allows access to the same reference data through Galaxy as well as e.g. command line, enabling reproducible analysis beyond the Galaxy community. Additional optimizations, e.g. using HTTP proxies or alien cache, are easily deployable across data centers, HPC clusters, and local labs, greatly facilitating scalability.

## Conclusions

We have extended the Galaxy platform to support the use of externally managed reference datasets using refgenie. Galaxy is able to seamlessly merge refgenie-provided references with those specified within its native Data Manager framework. We have provided a Data Manager tool that enables Galaxy administrators to view and pull additional refgenie assets graphically or using an API. This allows administrators to easily populate a Galaxy instance with standardized references to built-in data, improving interoperability of analysis among the now widely extended network of public Galaxy servers (https://galaxyproject.org/use/). Galaxy administrators are able to use standard Data Managers to build additional reference artifacts based upon refgenie provided asset sources. While these reference datasets will be available to Galaxy users, there is not yet a way to push them back into refgenie. Furthermore, although refgenie contains its own system for building assets, due to the lack of several important features, such as a system for declaring dependency requirements, adding this functionality to the Data Manager framework is left as a future endeavor.

We are pleased to report that this integration between refgenie and Galaxy was achieved without the need for architectural modifications to either tool and hope to see other examples of such collaboration in the future. The availability of these reference datasets via CVMFS, refgenie, Galaxy Data Managers, and other standard web protocols provides a globally-accessible network of provenance-backed reference datasets that are useful, accessible, and interoperable to a diverse range of researchers.

## Declarations

## List of Abbreviations

API: Application programming interface
CVMFS: CernVM File System
FAIR: Findability, Accessibility, Interoperability, and Reusability
FUSE: Filesystem in Userspace
GUI: graphical user interface

## Ethics approval and consent to participate

Not applicable

## Consent for publication

Not applicable

## Contributions

All authors contributed, read, and approved the manuscript.

## Acknowledgements

The authors would like to thank the communities and contributors of refgenie and Galaxy for their support and for creating the environments that allowed us to build this collaboration. We would also like to thank all who support open source software, and those who generate and distribute datasets following the FAIR principles.

