## supplementary materials for "Expanding the Galaxy’s reference data"

### refgenie for Local Instances

Built-in data files are critical for many Galaxy tools. This page will describe how to use **refgenie** with your local instance of Galaxy.

Please note that "built-in" or "cached" data can also be managed natively from within the Galaxy admin interface. For details, see: **Data Managers Overview** and our **Data Managers Tutorial**.

#### How it works

There are several steps needed for using refgenie with Galaxy. The first is to initialize a refgenie installation locally, following the standard directions at the **refgenie** website. Next, the galaxy.yml file needs to modified to point to the location of the `genomes.yml` file. Finally, an optional Data Manager tool can be installed from the ToolShed to enable a Galaxy administrator to populate the configured refgenie installation through the Galaxy interface.

##### Inititalize refgenie

Install refgenie and run `refgenie init`, e.g.:

```
/$ mkdir refgenie
/$ cd refgenie
/refgenie$ virtualenv -p python3 venv
  ...
/refgenie$ source venv/bin/activate
(venv) /refgenie$ pip install refgenie
  ...
(venv) /refgenie$ refgenie init -c genome_config.yaml
Initialized genome configuration file: /refgenie/genome_config.yaml
```

List available remote genomes:

```
(venv) /refgenie$ refgenie listr -c genome_config.yaml
  ...
```

Install a genome:

```
(venv) /refgenie$ refgenie pull -c genome_config.yaml -g t7 fasta
  ...
```

##### Configure Galaxy to load refgenie genomes

Edit `/$GALAXY_ROOT/config/galaxy.yml` to point to the refgenie genome configuration YAML file:

```
  # File containing refgenie configuration, e.g.
  # /path/to/genome_config.yaml. Can be used by refgenie backed tool
  # data tables.
  refgenie_config_file: /refgenie/genome_config.yaml
```

and then restart the Galaxy server.

##### Install a Data Manager tool for refgenie (optional)

For information on installing tools from the ToolShed, follow these directions. Search for "refgenie", to find the relevant tool.

To access and run Data Manager tools, follow these directions.

#### Figure 1. Extending Galaxy's reference data with refgenie.

**refgenie for Local Instances**
**How it works**
Inititalize refgenie
Configure Galaxy to load refgenie genomes
Install a Data Manager tool for refgenie (optional)
**Figure 1. Extending Galaxy's reference data with refgenie.**
**Table S1. Comparison of Galaxy, refgenie, and refgenie+Galaxy features**

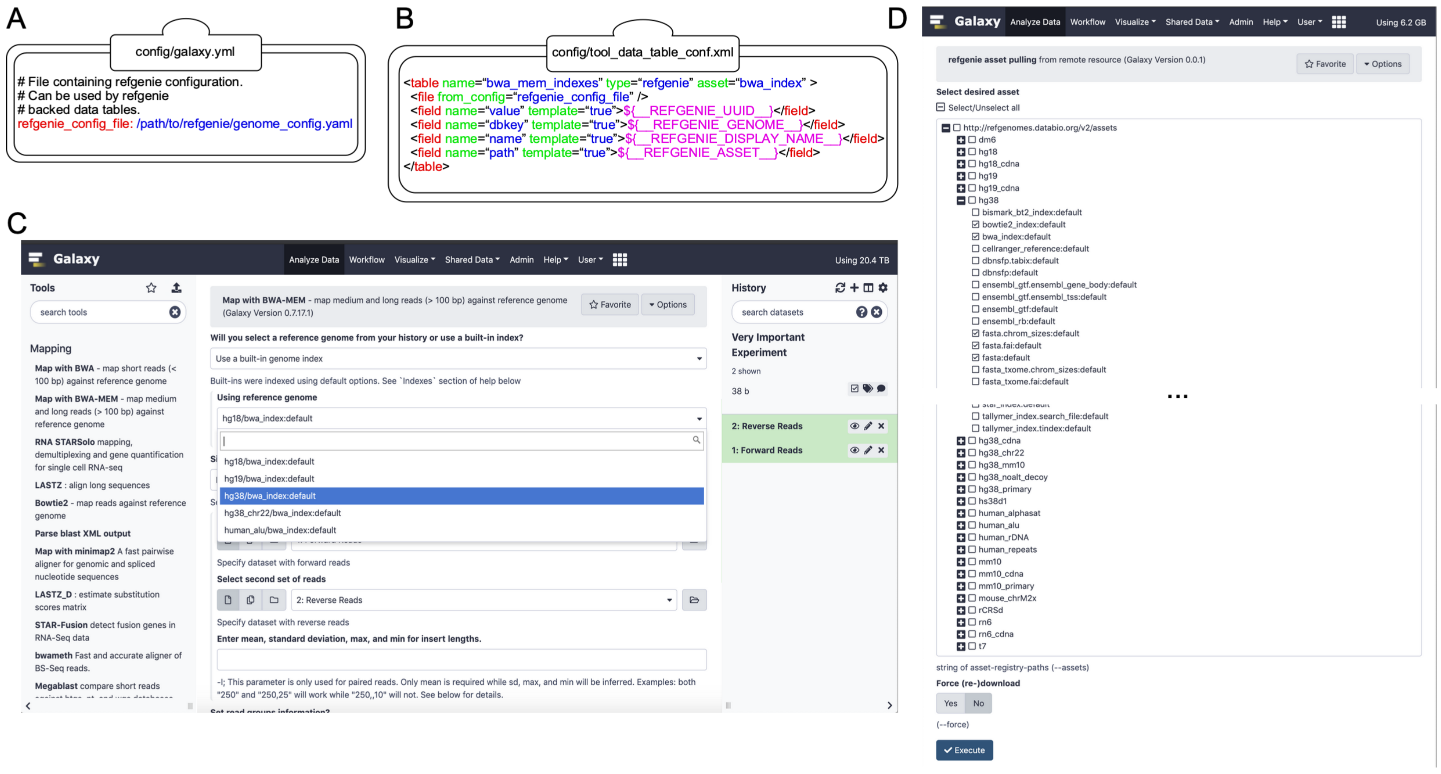


**Figure 1. Extending Galaxy's reference data with refgenie.** (A) Setting the value of 'refgenie_config_file' to the previously chosen genome configuration file path within the primary Galaxy configuration file (e.g. 'galaxy.yml'). (B) Example data table mapping between refgenie assets and Galaxy data tables for the BWA tool. Cheetah templating language is used to specify mappings between values, with several pre-populated refgenie variables available as shown. (C) refgenie assets are available for users to select and use in the Galaxy BWA tool. In this example, the user is mapping a set of paired-end sequencing reads against the hg38 genome. (D) A dynamically generated list of available remote refgenie assets are listed for an administrator to select in the 'refgenie pull' Galaxy Data Manager tool.

### Table S1. Comparison of Galaxy, refgenie, and refgenie+Galaxy features

| | refgenie | Galaxy | refgenie+Galaxy |
|---|---|---|---|
| features that refgenie has, but Galaxy does not have | X | | X |
| features that Galaxy has, but refgenie does not have | | X | X |

**Table S1. Comparison of Galaxy, refgenie, and refgenie+Galaxy features.** As indicated, refgenie+Galaxy contains the union of features of Galaxy and refgenie.
